# Embryonic Tissues as Active Foams

**DOI:** 10.1101/2020.06.17.157909

**Authors:** Sangwoo Kim, Marie Pochitaloff, Georgina-Stooke-Vaughan, Otger Campàs

## Abstract

The physical state of embryonic tissues emerges from non-equilibrium, collective interactions among constituent cells. Cellular jamming, rigidity transitions and characteristics of glassy dynamics have all been observed in multicellular systems, but there is no unifying framework to describe all these behaviors. Here we develop a general computational framework that enables the description of embryonic tissue dynamics, accounting for the presence of extracellular spaces, complex cell shapes and tension fluctuations. In addition to previously reported rigidity transitions, we find a distinct rigidity transition governed by the magnitude of tension fluctuations. Our results indicate that tissues are maximally rigid at the structural transition between confluent and non-confluent states, with actively-generated tension fluctuations controlling stress relaxation and tissue fluidization. Comparing simulation results to experimental data, we show that tension fluctuations do control rigidity transitions in embryonic tissues, highlighting a key role of non-equilibrium tension dynamics in developmental processes.

---

Many essential processes in multicellular organisms, from organ formation to tissue homeostasis, require a tight control of the tissue physical state^1, 2^. While tissue mechanics and structure at supracellular scales emerge from the collective physical interactions among the constituent cells, their control occurs at cell and subcellular levels. Bridging these scales is essential to understand the physical nature of active (non-equilibrium) multicellular systems and to identify the processes that cells use to control the physical state of embryonic tissues.

*In vitro* experiments of cell monolayers on substrates have revealed characteristics of glassy dynamics^3, 4^ and rigidity transitions^5–7^, which are thought to be linked to biological function and multiple pathologies. In contrast, suspended epithelial monolayers are largely solid-like *in vitro* ^8^ and show evidence of fracture *in vivo* ^9^. Experiments in embryonic tissues have shown characteristics of glassy dynamics in cell movements ^10^, viscous behavior at long-timescales ^11^ and also structural signatures reminiscent of jamming transitions ^12^. Recent *in vivo* experiments in developing zebrafish embryos showed the existence of a rigidity (fluid-to-solid) transition underlying the formation of the vertebrate body axis, revealing a functional role of rigidity transitions in embryonic development ^13^. It is, however, unclear how cells control rigidity transitions in multicellular systems and, more generally, whether all these observed phenomena share a common physical origin.

The physical behavior of multicellular systems has been studied theoretically using various approaches. Vertex models^14–19^ and Cellular Potts models^20, 21^, which account for cell geometry and use equilibrium formulations to describe the physical state of the system, predict a density-independent rigidity transition in confluent systems that is solely controlled by cell shape^15^. In contrast, self-propelled particle models, largely inspired in experiments of cultured cells that apply traction forces to move on synthetic substrates, account for the dynamics of the system and predict rigidity transitions that depend on cell density and self-propulsion^22, 23^. However, in most embryonic tissues (Fig. 1**a,b**), cells rarely apply net traction forces on substrates, but instead control tensions at the cell cortex, generating active force dipoles at cell-cell contacts. Recent observations have shown that both the presence of spaces between cells (non-confluence; Fig. 1**b,c**) and the dynamics of cell-cell contacts driven by active tensions (Fig. 1**e,f**) affect the physical (fluid/solid) state of the tissue ^13^.

**Figure 1:**
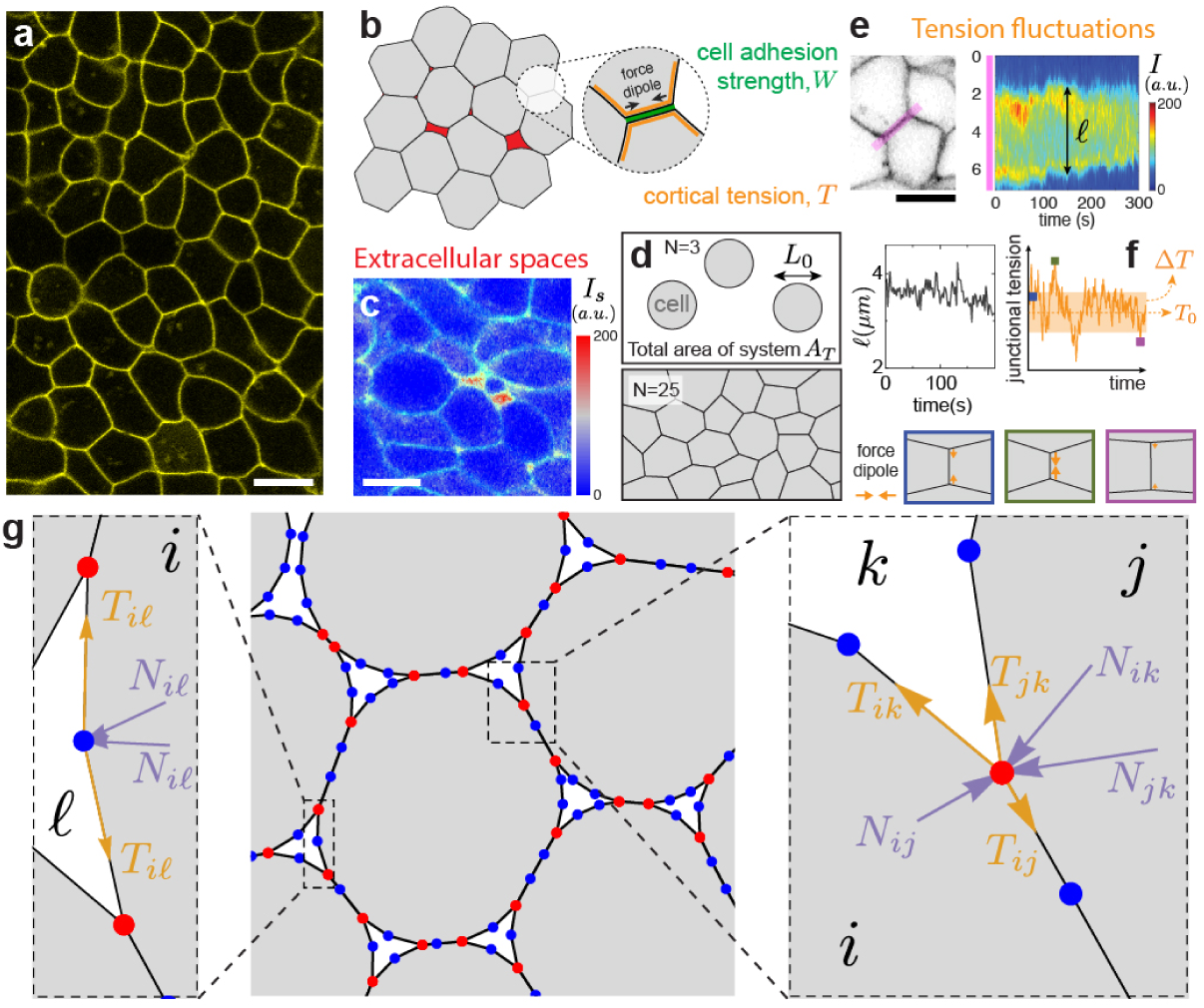
Characteristics of multicellular systems and simulation framework. **a**, Confocal section of embryonic tissue in a zebrafish embryo (membranes labelled; yellow). **b**, Schematics of cortical tension and adhesion at cell-cell contacts in a multicellular system with spaces (red) between cells (gray). **c**, Confocal section through zebrafish embryonic tissues showing the intensity (*I*_*s*_) of fluorescent reporter protein secreted extracellularly as well as cell membranes (green), indicating the presence of extracellular spaces (red range; Methods). **d**, Schematics defining the cell size *L*_0_, cell number *N* and simulation box area *A*_*T*_, which specify cell density. **e**, Kymograph of membrane signal intensity (*I*) along a tissue region (pink; left panel) containing a cell-cell contact, showing contact length *ℓ* fluctuations (bottom). **f**, Simulated tension fluctuations causing cell-cell contact length fluctuations. Increasing (decreasing) tension shortens (lengthens) cell-cell junctions (bottom). **g**, Schematics of the dynamic vertex model formulation. Triple vertices (physical vertices; red) and non-physical intermediate vertices (blue) are shown.

Here we develop a computational framework to study the dynamics, structure and physical state of active multicellular systems, including embryonic tissues. The simulations consist of a fully dynamic vertex model that accounts for extracellular spaces, complex cell shapes and the dynamics of tensions at cell-cell contacts. We recapitulate known density-dependent and density-independent rigidity transitions, and show the existence of a qualitatively distinct rigidity transition controlled by the magnitude of active tension fluctuations that occurs both in confluent and non-confluent systems. Comparing these simulations to quantitative data, our results indicate that the observed rigidity transition guiding vertebrate axis elongation ^13^ is mainly controlled by tension fluctuations.

## Dynamic Vertex Model with Extracellular Spaces

To study the dynamics of embryonic tissues, we generalize 2D vertex models by accounting for (i) extracellular spaces (treated as ‘cells’ with different physical characteristics; Fig. 1**b,c,g**; Methods), (ii) the stochastic dynamics of active cortical tensions (Fig. 1**f**) and (iii) complex cell shapes (implemented through addition of non-physical intermediate vertices; Fig. 1**g**; Methods). Tissue dynamics and structure, as well as cell movements and their shapes, are all determined by the dynamics of vertices (Fig. 1**g**), which follow from force balance, namely

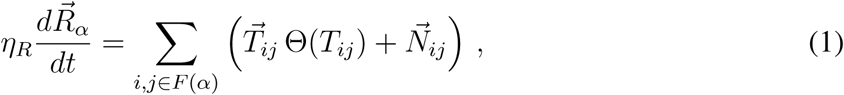

where 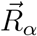 is the position of the vertex *α, η*_*R*_ is a friction coefficient characterizing the dissipation associated with moving a vertex, 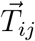 is the effective tension at the contact between cell *i* and *j* (with *F* (*α*) representing the set of all cells sharing vertex *α*) and 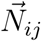 are the normal forces acting on vertex *α*. Finally, Θ(·) represents the Heaviside step function and prevents unrealistic negative tensions ^13, 24^.

Normal forces arise from osmotic pressure differences in adjacent cells, namely *N*_*ij*_ = (Δ*P*_*i*_ − Δ*P*_*j*_) *L*_*ij*_/2, where *L*_*ij*_ and Δ*P*_*i*_ (Δ*P*_*j*_) are, respectively, the contour length of the contact between cells *i* and *j* and the osmotic pressure difference across cell *i* (*j*). While we use the experimentally measured relation between osmotic pressure and cell volume ^25^ (area in 2D), the specific functional form does not qualitatively affect our results as long as cell volume decreases with increasing osmotic pressure (Methods).

To capture key observed features of the dynamics of effective tensions, *T*_*ij*_, namely their fluctuating nature and their finite persistence time ^13^, in a simple, yet generic manner, we write them as an Ornstein-Uhlenbeck process with a tension that fluctuates around a fixed point 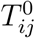 and has a persistence time *τ*_*T*_ ^26, 27^, specifically

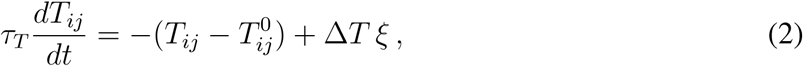

where Δ*T* is the amplitude of tension fluctuations and *ξ* is gaussian white noise (Fig. 1**f**; Methods). The fixed point effective tensions depend on both the average cortical tension, *T*_0_, and average strength of cell-cell adhesion, *W*, which, like Δ*T*, are different at cell-cell contacts and free cell boundaries (Fig. 1**b**; Methods).

Scaling all quantities with characteristic scales, we obtain a dimensionless set of equations as well as the relevant dimensionless parameters (Methods). The dimensionless parameter Δ*T/T*_0_ characterizes the magnitude of tension fluctuations (Fig. 1**e,f**), the ratio *W/T*_0_ characterizes the relative strength of cell-cell adhesion and cortical tension (Fig. 1**b**), the parameter *P*_0_*L*_0_*/T*_0_ measures the relative magnitude of normal to tensional forces, the ratio *τ*_*T*_ */τ*_*R*_ of the persistence time of tension fluctuations and the characteristic timescale *τ*_*R*_ ≡ *η*_*T*_ *L*_0_*/T*_0_ of dissipation at vertices and, finally, the normalized system density 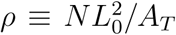, with *A*_*T*_ and *N* being the total area of the system and the number of cells, respectively (Fig. 1**d**). Since *P*_0_*L*_0_*/T*_0_ and *τ*_*T*_ */τ*_*R*_ can be estimated from existing experimental data (Methods), we focus on the parameter space spanned by Δ*T/T*_0_, *W/T*_0_ and *ρ*.

## Structural Transitions at Equilibrium

In the absence of tension fluctuations (Δ*T/T*_0_ = 0), the amount of extracellular spaces in equilibrium configurations is determined by force balance and varies with both the cell density *ρ* and the relative cell adhesion strength *W/T*_0_ (Fig. 2**a**). Increasing cell density results in larger cellular volume fraction *ϕ* (Fig. 2**b**) and cell contact number *z* (Fig. 2**c**), with the system eventually becoming confluent (*ϕ* = 1) at an adhesion-dependent critical density, *ρ*_*c*_(*W/T*_0_). For any fixed cell density, the system undergoes a non-confluent to confluent transition as *W/T*_0_ is increased, since higher adhesion promotes stronger cell-cell contacts (Fig. 2**a,b**).

**Figure 2:**
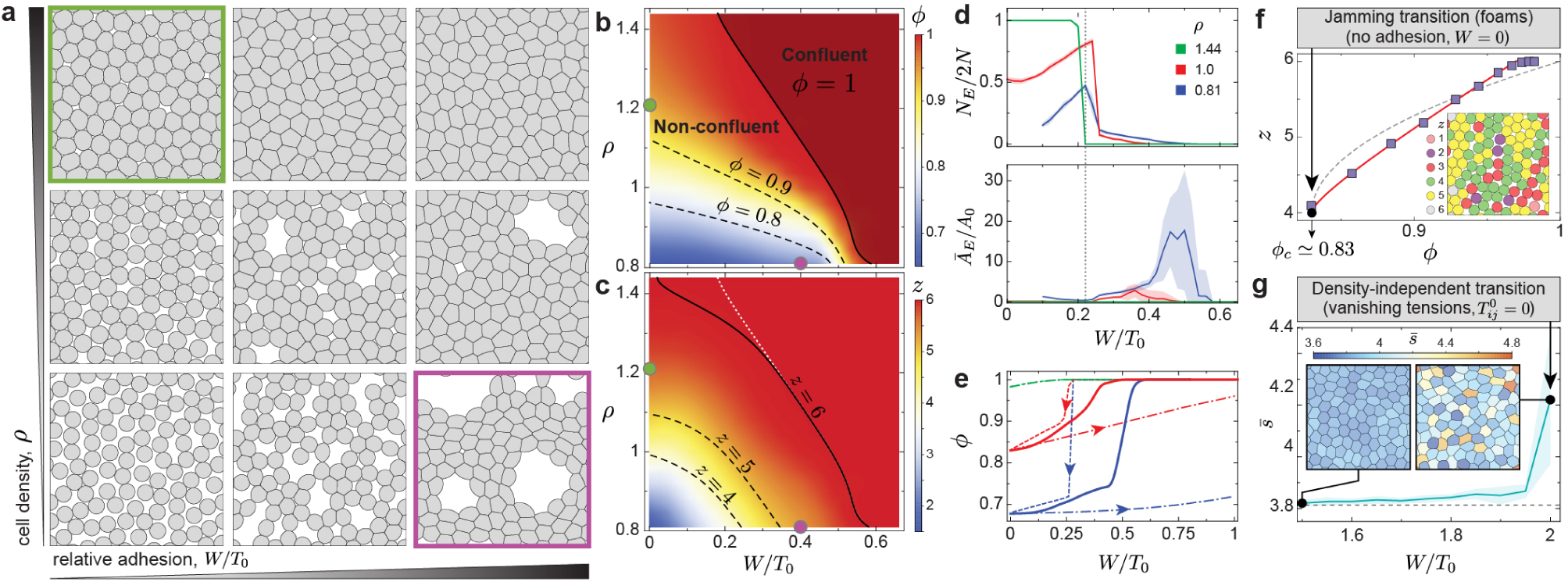
Equilibrium configurations and structural transitions. **a-c**, Representative equilibrium configurations (a), volume fraction *ϕ* (b) and mean number of neighbors *z* (c) for varying values of relative adhesion *W/T*_0_ and cell density *ρ*. **d**, number *N*_*E*_ (top) and average area *Ā*_*E*_ (bottom) of extracellular spaces for varying relative cell adhesion and different cell densities, showing a sharp structural transition at *W/T*_0_ *≈* 0.23 (gray line) leading to the opening of large extracellular spaces. **e**, Lowering (dashed line) and increasing (dash-dotted line) relative adhesion quasi-statically shows bistable states and strong hysteresis in equilibrium configurations. Equilibrium quenched states are also shown (solid line; Methods). **f**, Average neighbor number (cell contacts) *z* as the system volume fraction changes at vanish cell adhesion, showing the existence of a jamming transition at *ϕ*_*c*_ ≃ 0.83 (configuration shown in inset). Power law fits *z* − *z*_*c*_ = *z*_0_(*ϕ* − *ϕ*_*c*_)^1/2^ + *z*_1_(*ϕ* − *ϕ*_*c*_) with non-zero *z*_0_ and *z*_1_ (red line) and with *z*_1_ = 0 (gray line) are shown. **g**, Average shape factor for varying relative adhesion showing a sharp increase at *W/T*_0_ = 2 (vanishing tensions), leading to anisotropic cell shapes (inset), recapitulating density-independent transitions. Error bands = SD. N= 20 (b,c,d,f,g) and N=10 (e) independent simulations for each parameter set.

Changes in relative cell adhesion not only affect the volume fraction but also the structural characteristics of extracellular spaces. Equilibrium configurations sharply transition from a large number of small extracellular spaces to a few large extracellular spaces at *W/T*_0_ *≈* 0.23 (Fig. 2**a,d**), as small triangular extracellular spaces can only be stabilized below this value (Supplementary Information). Concomitant to the presence of large extracellular holes, the spontaneous clustering of cells strongly resembles flocculation in sticky emulsions ^28, 29^ (Fig. 2**a**). Moreover, the system displays bistability between two possible equilibrium configurations for *W/T*_0_ ≥ 0.23, namely a confluent state with stretched cells and a non-confluent state with sparse and large extracellular spaces (Fig. 2**e**). While the most stable branch switches from a non-confluent state with a few large extracellular spaces to the confluent state as *W/T*_0_ increases, changing adhesion (or cortical tension) in a quasi-static manner leads to strong hysteresis in equilibrium configurations (Fig. 2**e** and Supplementary movie 1; Methods).

In the limit of vanishing cell adhesion, the system should behave as foams/emulsions, which display a jamming transition at a critical value *ϕ*_*c*_. The isostatic condition (*z* = 4 in 2D) sets the critical volume fraction *ϕ*_*c*_ *≈* 0.83 of the system^30^ (Fig. 2**f**). Both this *ϕ*_*c*_ value and the power-law dependence of *z* (Fig. 2**f**), are in agreement with recent equilibrium simulations of deformable particles ^31^, indicating that our description accurately describes the foam limit.

Previously reported density-independent rigidity transitions in confluent states are also captured within this framework. In these transitions, which are controlled by the cell shape factor *s* (defined as 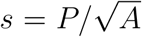, with *P* and *A* being the cell perimeter and area, respectively), the system switches from a solid to a fluid state at *s*_*c*_ ≃ 3.81, with cells transiting from isotropic to anisotropic shapes ^15^. Importantly, the fluid state in these descriptions is characterized by vanishing effective tensions 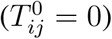, allowing neighbor exchanges at no energetic cost ^17^. In the present framework, density-independent transitions are recapitulated at *W/T*_0_ = 2, which leads to vanishing effective tensions and associated fluid state, as well as a sharply larger average shape factor due to the emergence of anisotropic cell shapes (Fig. 2**g**).

## Dynamic Simulations with Finite Tension Fluctuations

In the presence of finite tension fluctuations (Δ*T/T*_0_ *>* 0), the tension dynamics at cell-cell contacts can drive cell movements, neighbor exchanges and cell shape changes (Supplementary movies 2-3). At timescales longer than all characteristic timescales (*t ≫ τ*_*T*_ *> τ*_*R*_), cell movements are caged for small Δ*T/T*_0_, as indicated by the saturation of the Mean Squared Displacement (MSD) and bounded cell trajectories (Fig. 3**a,b**; Methods). For increasing magnitudes of tension fluctuations, cell uncaging starts to occur and the asymptotic behavior of the MSD for *t ≫ τ*_*T*_ becomes a power law (MSD ∼ *t*^*α*^), with an exponent *α* that increases with activity (Fig. 3**a** inset). This evidences sub-diffusive (*α <* 1) cell movements for intermediate activities, and diffusive (*α* = 1) behavior for large enough tension fluctuations (Fig. 3**a,b**).

**Figure 3:**
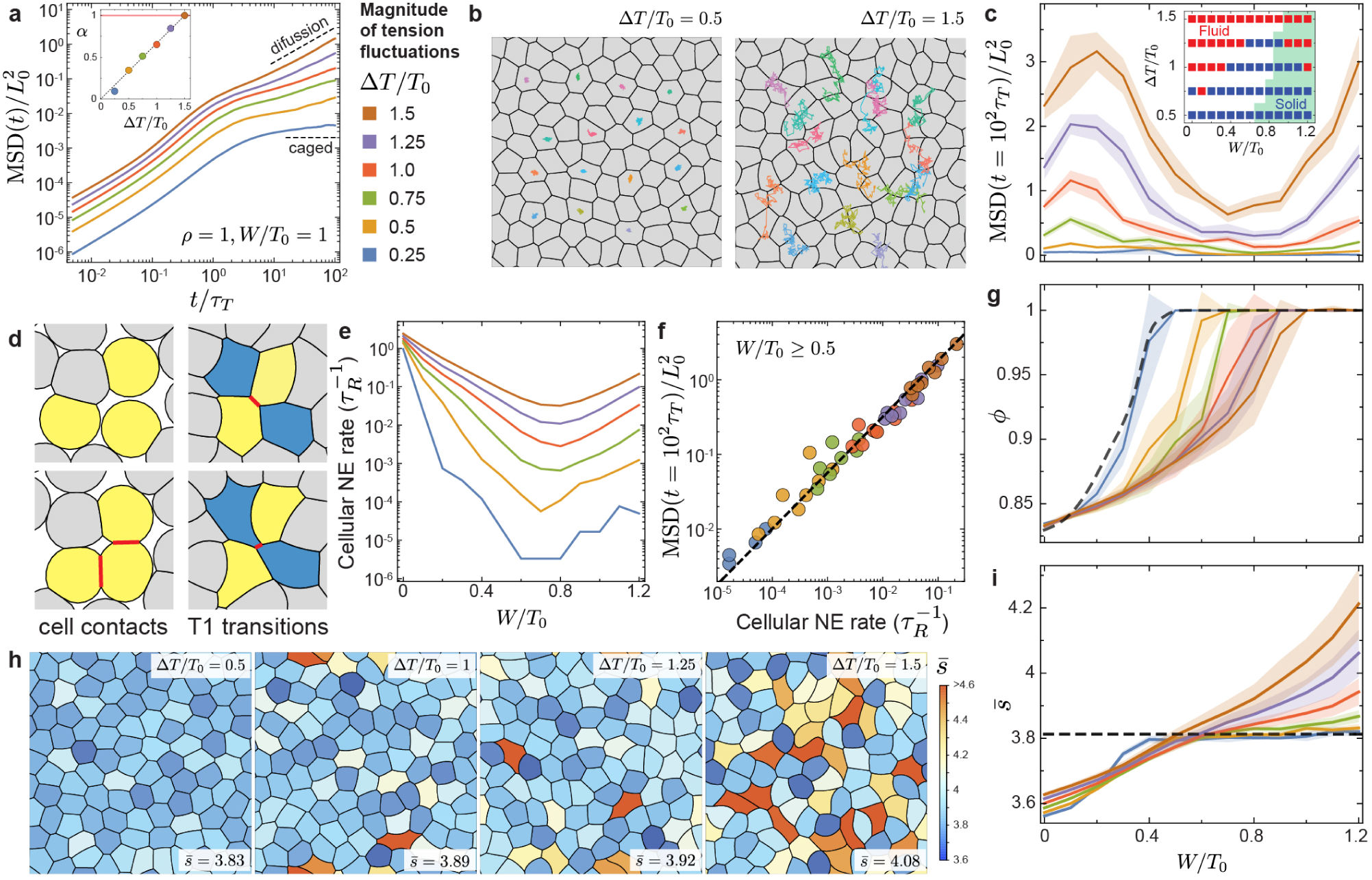
Tissue dynamics and structure with finite tension fluctuations. **a**, MSD for varying magnitudes of tension fluctuations Δ*T/T*_0_, showing sub-diffusive (0 *< α <* 1) and diffusive (*α* = 1) behaviors as tension fluctuation increase (inset). **b**, Snapshots of dynamic configurations with examples of cell trajectories over *t/τ*_*T*_ = 10^2^. **c**, MSD at long timescales (*t* = 10^2^*τ*_*T*_ *≫ τ*_*T*_ *> τ*_*R*_), showing non-monotonous behavior for varying relative adhesion strength and minimal values at the structural transition. Qualitative solid-fluid phase diagram obtained from the characteristics of cell movements (inset). **d**, Distinct types of neighbor exchange events: gain/loss of cell contacts (left) and T1 transitions (right). **e**, Cellular neighbor exchange (NE) rate for varying relative adhesion. **f**, Power law relation (with exponent 3/4) between the long timescales MSD and the cellular NE rate. **g-i**, Dependence of the system volume fraction *ϕ* (g) and average shape factor 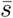 (i) on relative adhesion for different magnitudes of tension fluctuations. Representative snapshots of dynamic configurations for increasing tension fluctuations are shown (h). Error bands = SD. N=10 independent simulations for each parameter set.

Comparing MSD values at long timescales (*t* = 10^2^*τ*_*T*_ *≫ τ*_*T*_ *> τ*_*R*_) shows that increasing tension fluctuations always leads to higher MSD values, regardless of cell adhesion strength. However, for a fixed level of activity, MSD values vary non-monotonically as adhesion increases, displaying very reduced cell movements at the transition between non-confluent to confluent states (Fig. 3**c**). Associating MSD values above (below) half the cell size with fluid (solid) states (Methods), it is possible to qualitatively assess the physical state of the system (Fig. 3**c**, inset). Irrespective of the adhesion strength, the system remains solid for small tension fluctuations and becomes fluid for strong enough tension fluctuations. However, for any given intermediate level of activity, there are fluid phases at both low and high adhesion levels, separated by a solid phase for intermediate adhesion, with the system being most solid-like (cell movements most restricted) at the non-confluent to confluent transition. Increasing adhesion in a non-confluent system leads to higher volume fraction, making it more solid-like. In contrast, increasing adhesion in a confluent system leads to lower energy barriers for neighbor exchange and a more fluid-like system.

Both the system’s rigidity and uncaging events are directly related to changes in cell contact topology (neighbor exchanges). While generation and loss of cell contacts dominate in the non-confluent regime, neighbor exchanges shift progressively to T1 transitions as the system approaches confluence for higher adhesion levels (Fig. 3**d,e**). After reaching confluence, tension fluctuations induce T1 transitions that allow cells to escape the neighbor-imposed cages and MSD values scale with the neighbor exchange rate, *k*_NE_, as 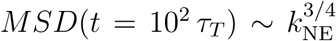 (Fig. 3**f**). The timescale for T1 transitions, 1*/k*_NE_, diverges as the activity vanishes (or *k*_NE_ → 0 as Δ*T/T*_0_ → 0), a signature of glassy dynamics.

Beyond dynamics, the structural features of the system also depend on activity. Tension fluctuations generally promote transitions from confluent to non-confluent states by opening up extracellular spaces at weak regions (Fig. 3**g**), implying that larger adhesion values are required to reach confluent states for increasing Δ*T/T*_0_. This effect becomes negligible for large cell densities (*ρ >* 1) and volume fraction is then solely determined by cell density *ρ* and relative cell adhesion *W/T*_0_ (Supplementary Information). While cell shapes are only moderately affected by tension fluctuations in non-confluent regimes, with low shape factors associated with rounder cells, increasing tension fluctuations in confluent regimes leads to substantially larger shape factors (Fig. 3**h,i**). Despite sharing similar geometric features, the rigidity transition induced by tension fluctuations is qualitatively different from the density-independent transition, which requires junctional tensions to vanish. Here, cell-cell contacts maintain finite tensions (Supplementary Information; Supplementary movie 3) and it is spatiotemporal tension fluctuations that generate large shape factors in the confluent regime (Fig. 2**g**).

## Physical state of multicellular systems

To directly assess the physical state of the system, we monitor shear stress relaxation after imposing an affine (shear) deformation (Fig. 4**a**; Methods). Since tissue fluidization in biological tissues is associated with the non-linear mechanical response ^13^, we impose large strains *ϵ*_*xy*_ of 150% (*ϵ*_*xy*_ = 1.5). In the absence of tension fluctuations, the initial stress jump is largest with no adhesion (Fig. 4**b**), and vanishes when effective tensions vanish (*W/T*_0_ = 2; Supplementary Information). Subsequently, shear stress relaxes with a characteristic timescale *τ*_*R*_ towards a constant value at long timescales, namely the yield stress *σ*_*Y*_ ^32^, which depends non-monotonically on cell adhesion (Fig. 4**c**; Supplementary movie 5). For low relative adhesion *W/T*_0_, the system is non-confluent and the yield stress increases with increasing adhesion as extracellular spaces close down. In contrast, for adhesion values leading to confluence, increasing *W/T*_0_ leads to a decreasing yield stress due to lower effective tensions. The vanishing yield stress at *W/T*_0_ = 2 indicates a fluid state, in agreement with density-independent rigidity transitions (Fig. 2**g**). At equilibrium, the system is maximally rigid (maximal yield stress) at the structural transition between confluent and non-confluent states (Fig. 4**c,d**).

**Figure 4:**
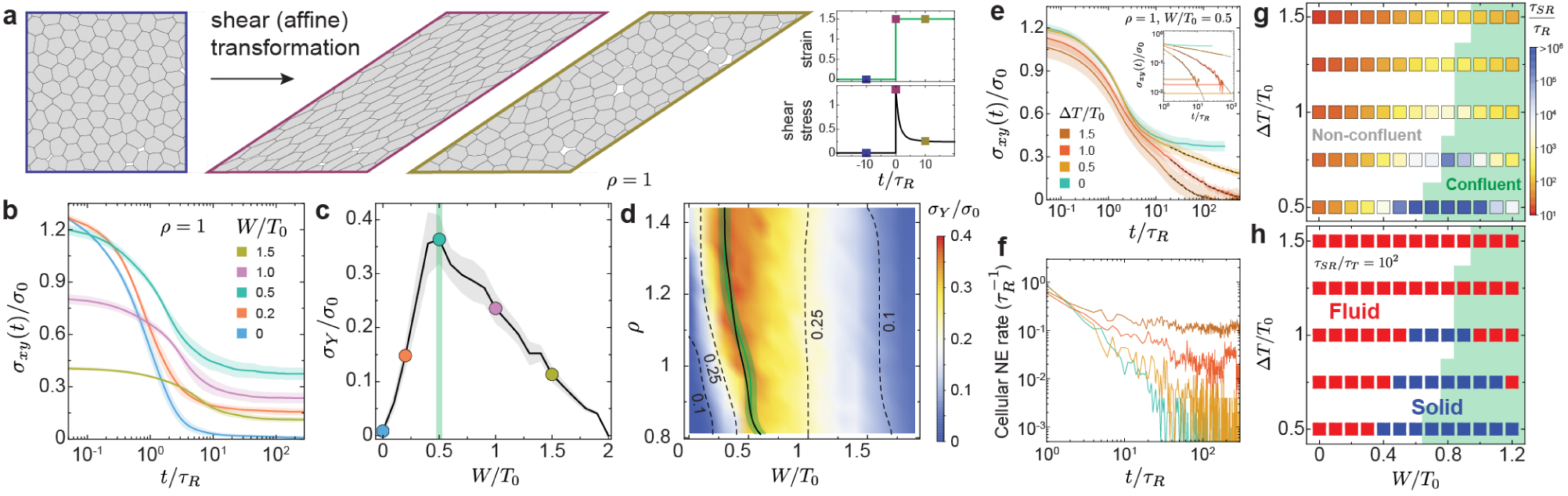
Stress relaxation in both passive and active multicellular systems. **a**, Schematics of a simple shear deformation imposing a large strain step (*ϵ*_*xy*_ = 1.5), with associated temporal evolution of both strain and shear stress. **b**, Temporal relaxation of shear stress *σ*_*xy*_ (normalized to 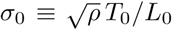) after the imposed strain step for varying adhesion levels and in the absence of tension fluctuations. **c-d**, Dependence of the yield stress *σ*_*Y*_ on the relative adhesion strength (c) and on both cell density and relative adhesion (d), showing a maximum at the structural transition between confluent and non-confluent states (green line). **e-f**, Stress relaxation (e) and temporal changes in cellular NE rate (f) after the imposed strain step for varying magnitudes of tension fluctuations. The long timescale stress relaxation follows stretched exponentials (black dashed lines) and eventually reaches the average value *σ*_*A*_ of active shear stress fluctuations (horizontal lines in inset). The cellular NE rate quickly decays to zero in the absence of activity, but remains finite in the presence of activity, with actively-driven neighbor exchanges enabling further stress relaxation. **g**, Stress relaxation timescale *τ*_*SR*_ for varying magnitude of tension fluctuations and relative adhesion, showing a sharp increase as the structural transition between non-confluent and confluent (green background) states. **h**, Phase diagram showing the transition between fluid and solid tissue states for different activity values and relative adhesion strength. Solid states surround the structural transition and are found both in confluent and non-confluent configurations for low enough activity. Error bands = SD. N=10 independent simulations for each parameter set.

Introducing finite tension fluctuations qualitatively changes stress relaxation at long timescales, which displays a slow stress decay rather than plateauing (Fig. 4**e**; Supplementary movie 6). This long timescale stress relaxation is driven by actively-induced T1 transitions (Fig. 4**f**), and can be accurately described by a stretched-exponential function (Fig. 4**e** inset), as previously done to explain the dynamics of systems with a large number of intrinsic timescales ^33, 34^. The timescale of stress relaxation, *τ*_*SR*_, at which shear stress reaches the level of active shear stress in the unperturbed system (Fig. 4**e** inset; Supplementary Information), varies by over five orders of magnitude as the magnitude of tension fluctuations or relative adhesion change slightly (Fig. 4**g**). While larger activity values reduce *τ*_*SR*_ monotonically, increasing the relative adhesion strength leads to drastic and non-monotonic changes in *τ*_*SR*_, which rapidly increases in non-confluent states and displays the opposite behavior in confluent states. The largest stress relaxation timescale occurs at the structural transition between non-confluent to confluent states, in agreement with our previous results (Fig. 3**c,d** and Fig. 4**b**).

As in colloidal glasses^35^, whether the system behaves as a fluid or a solid depends on observation timescales. In embryonic tissues, if the characteristic timescale *τ*_*d*_ of developmental processes is larger (smaller) than the stress relaxation timescale *τ*_*SR*_, namely *τ*_*d*_*/τ*_*SR*_ *≫* 1 (*τ*_*d*_*/τ*_*SR*_ *≪*1), the tissue behaves as a fluid (solid). Using typical developmental timescales (*τ*_*d*_ ∼ 1-2 hours; *τ*_*d*_*/τ*_*T*_ ∼ 10^2^), we obtain the tissue phase diagram (Fig. 4**h**). Over a critical value of tension fluctuations, the tissue is always fluid regardless of cell adhesion levels. Below that critical activity value, the tissue is fluid at both low and high adhesion levels, but solid in between, in the region of the phase diagram surrounding the structural transition between non-confluent to confluent states.

## Rigidity transition during vertebrate axis elongation

Recent experiments have shown that during body axis elongation in zebrafish, posterior tissues transit from a fluid state in the mesodermal progenitor zone (MPZ) to a solid state in the presomitic mesoderm (PSM) (Fig. 5**a,b**). To study the nature of rigidity transitions in these embryonic tissues, we compare our simulation results to both new and existing data ^13^ of the structure, dynamics and mechanics of the MPZ and PSM tissues (Fig. 5).

**Figure 5:**
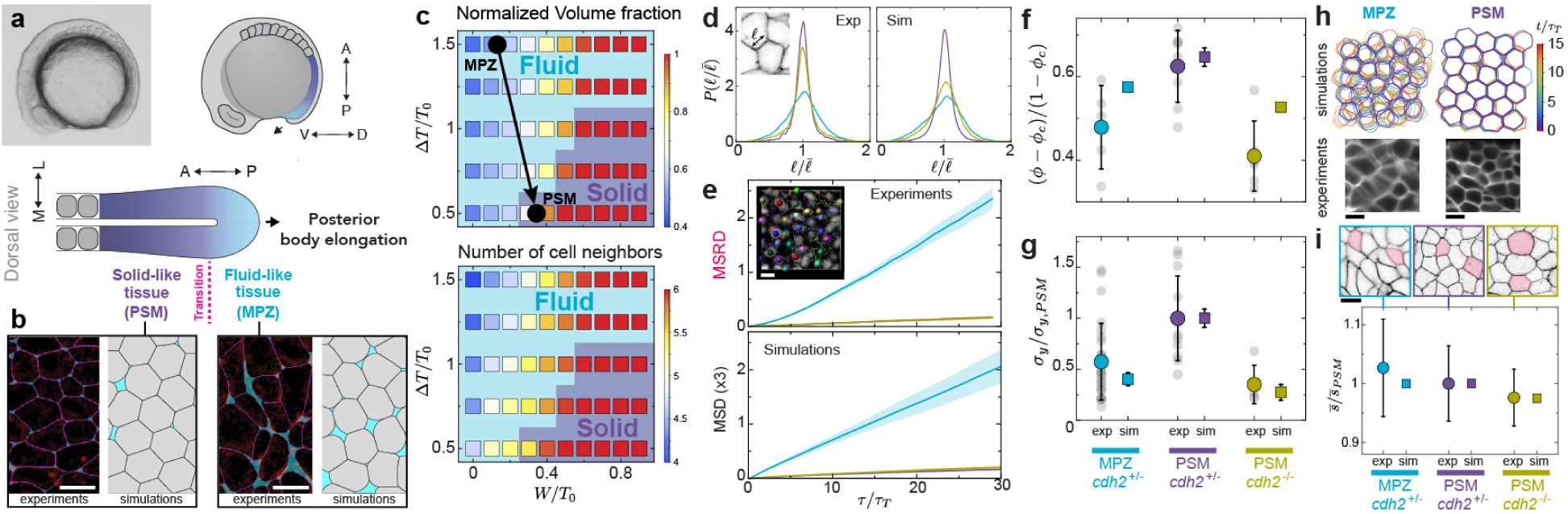
Nature of rigidity transitions in embryonic tissues. **a**, Lateral view of a zebrafish embryo (a, top left) and sketches showing a lateral view (a, top right) and a dorsal view of posterior tissues (bottom) indicating the transition between a fluid tissue state in the MPZ (cyan) and a solid state in the PSM (violet). **b**, Confocal sections of the MPZ and PSM tissues in a 10-somites stage zebrafish embryo (Methods), showing segmented cell borders (pink) and extracellular spaces (light blue). Simulated tissue configurations (cells in gray) are also shown. **c**, Phase diagram showing fluid and solid tissues states (background) for varying relative adhesion and magnitude of tension fluctuations, with the corresponding values of normalized volume fraction (*ϕ* − *ϕ*_*c*_)/(1 − *ϕ*_*c*_) (top) and mean neighbor number *z* (bottom) overimposed. The parameter values that best fit the experimental data for MPZ and PSM are shown (black dots), and the arrow connecting them represents the transition between fluid to solid states. **d-g**, Comparison of experimental (exp) data and simulations (sim) for MPZ (cyan) and PSM (violet) tissues in zebrafish embryos (*cdh2*^+/−^; wild type phenotype), as well as in the PSM of N-cadherin mutants (green; *cdh2*^−/−^): distribution 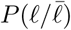 of cell-cell contact length fluctuations (d; inset shows a confocal section defining the cell-cell contact length *ℓ*), MSD and MSRD (e; Mean Squared Relative Displacement, or MSRD, is used for experimental data; MSRD corresponds to 3x MSD; inset shows nuclear detection and tracking; Methods), normalized volume fraction (*ϕ* − *ϕ*_*c*_)/(1 − *ϕ*_*c*_) (f), and yield stress *σ*_*Y*_ (normalized to the PSM value, *σ*_*Y,P SM*_). **h**, Experimentally measured and simulated dynamics of cell shapes in both MPZ and PSM (Methods), showing faster dynamics in the MPZ (blurred borders) and largely static cell boundaries in the PSM. Experimental data is an average intensity projection of a confocal section timelapse (membrane-labeled embryos; Methods). **i**, Average measured (exp) and simulated (sim) shape factor 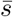 (normalized to PSM values,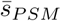) for the different cases, with confocal snapshots and representative cell shapes (pink; top panels) shown (PSM, n= 230; MPZ, n= 242; PSM *cdh2*^−/−^, n = 244. n= cells analyzed). Data in panels d-g is from reference ^13^, with the exception of the MSRD in the PSM of *cdh2*^−/−^ mutants (e; n=850 cells). Error bands = SD (e, sim) and SEM (e, exp); Error bars = SD elsewhere. N=10 independent simulations for each parameter set.

The presence of extracellular spaces in both the MPZ and PSM (Fig. 5**b**; Methods) restrict the values of the relative adhesion parameter to *W/T*_0_ ≲ 0.6 (Fig. 5**c**), with the cell density being the same in both regions ^13^. Simultaneously fitting the distribution of cell-cell contact length fluctuations (Fig. 5**d**), the measured MSD (Fig. 5**e**), the cell volume fraction (Fig. 5**f**) and the yield stress (Fig. 5**g**), it is possible to reproduce all observations, provided that tensional fluctuations and relative cell adhesion are larger in the MPZ than in the PSM (Δ*T/T*_0_ = 0.5, 1.5 and *W/T*_0_ = 0.3, 0.2 in PSM and MPZ, respectively). It is also possible to reproduce the observed higher cell boundary dynamics in the MPZ than in the PSM (Fig. 5**h**) and the relative changes in cell shape factor from PSM to MPZ (Fig. 5**i**), albeit not quantitatively due to the larger measurement error for cell shapes. For the density values that match experimental data (*ρ* = 1.1025 in MPZ and PSM), the observed differences in volume fraction between MPZ and PSM can only be explained by changes in relative adhesion from MPZ to PSM, but not by changes in the magnitude of tension fluctuations (Supplementary Information). This result indicates that during axis elongation, cells adjust their relative cell adhesion to control the level of extracellular spaces, differentially predisposing the MPZ tissue to cell uncaging. Tension fluctuations are then used to actually fluidize the tissue. Indeed, cell movements in the PSM of N-cadherin (*cdh2*^−/−^) mutants (Methods), which lack N-cadherin adhesion ^36^ and are characterized by reduced cell volume fraction, lower yield stress but similar cell-cell contact fluctuations as wild type embryos ^13^ (Fig. 5**d,f,g**), show the same MSD behavior as wild type embryos and a solid-like tissue state (Fig. 5**e**). Placing the obtained parameter values for MPZ and PSM in the phase diagram indicates that the magnitude of tension fluctuations plays a key role in the control of the observed fluid-to-solid transition (Fig. 5**c**).

## Discussion

We developed a general framework to study the structure, dynamics and mechanics of active (non-equilibrium) multicellular systems in a wide range of conditions. While jamming transitions and density-independent transitions are controlled by cell density and cell shape, respectively, we show that tension fluctuations control a distinct rigidity transition in both confluent and non-confluent states. These actively-generated tension fluctuations drive structural rearrangements that relax stresses, thereby fluidizing the tissue. Unlike density-independent rigidity transitions, all cell-cell contacts maintain finite tensions across the fluctuation-mediated rigidity transition, even when the tissue is in a fluid state.

Our results indicate that embryonic tissues are maximally rigid at the structural transition between confluent and non-confluent states. The existence of bistability between configurations having small spaces between cells and configurations with large extracellular ‘holes’, could lead to hysteretic behavior and discontinuous transitions between these states if cells in tissues slowly change control parameters (average cortical tension or adhesion) during development. While most embryonic tissues likely avoid this regime, as it could lead to epithelial fracture and loss of its barrier function, recently reported dynamics and fracture of epithelial monolayers in the organism *Trichoplax Adhaerens* display features of this structural transition ^9^.

Finally, quantitatively comparing our simulations to experimental data of embryonic tissues during vertebrate body axis elongation indicates that tension fluctuations play a prominent role in the control of the observed rigidity (fluid-solid) transition. At short timescales, before actively-induced T1 transitions occur, the tissues are solid-like and behave like wet foams, with adhesion levels controlling the degree of cellular confinement. However, long timescale stress relaxation and tissue fluidization is caused by actively-induced structural rearrangements controlled by tension fluctuations. These results highlight the relevance of tension fluctuations, rather than average cortical tension or adhesion, in the control of the tissue physical state and rigidity transitions in embryonic tissues, and suggest that cells may have mechanisms to specifically control the characteristics of tension fluctuations.

## Acknowledgements

We thank all members of the Campàs group for their comments and help, Payam Rowghanian for help with cell segmentation and the UCSB Animal Research Center for support. We also thank Holger Knaut (New York University) and Sean Megason (Harvard University) for kindly providing the Tg(hsp70:secP-mCherry)^p1^ and Tg(actb2:memCherry2)^hm29^ transgenic lines, respectively. The Tg(actb2:mem-neonGreen-neonGreen)^hm40^ line was generously provided before publication by Toru Kawanishi and Ian Swinburne in Sean Megason’s lab (Harvard University). This work was supported by the Eunice Kennedy Shriver National Institute of Child Health and Human Development of the National Institutes of Health (R01HD095797). We acknowledge support from the Center for Scientific Computing from the CNSI, MRL: an NSF MRSEC (DMR-1720256) and NSF CNS-1725797.

## Methods

### Pressure and volume relation

The (osmotic) pressure *P* of a cell has been experimentally shown to vary with its volume according to *P* = *K/V* ^25^, meaning that changes in the osmotic pressure difference between the inside (*P*) and outside (*P*_0_) of the cell lead to changes in the cell volume *V*, with *K* characterizing the cell compressibility (we assumed for simplicity that cells are never compressed close to their dry mass limit). In our 2D description, the cell area *A* plays the role of the volume, so that *P* = *K/A*, with the characteristic cell size *L*_0_ (or preferred area 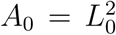) being set by this relation, namely 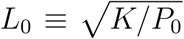. However, the specific P-V (or P-A) functional form does not change our results qualitatively as long as the cell volume decreases with increasing applied osmotic pressure, namely *P* − *P*_0_ = −Γ(*A* − *A*_0_) (with Γ being a coefficient characterizing the cell’s compressibility), and the deviations from the preferred cell area in 2D (*A*_0_) are mild.

### Fixed point tension and tension dynamics

The fixed point tension, 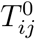, corresponds to an average effective tension at cell-cell contacts and has contributions from the average actomyosin-generated cortical tension *T*_0_ in each cell and the average cell-cell adhesion strength *W*, so that 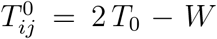 (Fig. 1**b**). At free cell boundaries (cell boundaries contacting the extracellular space), the average effective tension is just the average cortical tension of the cell, namely 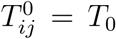. Here we assume the average tension *T*_0_ not to change with the cell perimeter, as no current experimental observations suggest that dependence. In situations where the cell shape becomes very anisotropic or contorted enough to have very large perimeters, a dependence of the average effective tension on the perimeter could potentially arise from limitations in plasma membrane availability.

The stochastic dynamics of tensions depends on the magnitude of tension fluctuations, which is Δ*T* at cell-cell contacts and Δ*T/*2 at free cell boundaries (cell boundaries contacting the extra-cellular space).

### Parameter estimation

Since the relaxation timescale *τ*_*R*_ has been measured to be much smaller than the persistence timescale of tension variations (*τ*_*R*_ *≲* 20s ^37, 38^; *τ*_*T*_ ≃ 90s ^13^) we set the ratio to *τ*_*T*_ */τ*_*R*_ = 10. While the values of osmotic pressure are unknown *in vivo*, they are expected to be larger than cortical stresses ^25^, so that *P*_0_*L*_0_*/T*_0_ *≫* 1. Consequently, we fix *P*_0_*L*_0_*/T*_0_ = 10 for all simulations. This ensures relatively mild cell size variations, as observed experimentally ^13^. Fixing these parameters reduces the parameter space to the normalized amplitude of tension fluctuations Δ*T/T*_0_, the ratio *W/T*_0_ of average adhesion strength and average cortical tension, and the ratio 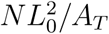 of the cell’s total preferred area and the total available area.

### Numerical integration

The dimensionless version of the governing equations (Eqs. 1-2; Supplementary Information) was integrated numerically using Euler-Maruyama method with a time step, Δ*t*. We used Δ*t* = 0.005 *τ*_*R*_ for all simulations to have a time resolution much smaller than the smallest characteristic timescale in the system, *τ*_*R*_.

### Initial configuration generation protocol

A polygonal tiling of confluent states is first generated by random Poisson Voronoi tessellation in a square periodic box of total area *A*_*T*_. Prior to the introduction of extracellular spaces, the initial confluent configuration is first annealed to a local equilibrium state to prevent sudden adjustment of cell shape with extracellular spaces. Extracellular spaces are introduced by replacing each vertex in the confluent configurations with a small triangular extracellular space ‘cell’ centered around the original vertex position, and with the 3 new vertices located on the each of the original edges and of 1% their original length. Small tension fluctuations (Δ*T/T*_0_ = 0.5) are applied for a duration *τ*_*T*_. The resulting configuration is used as an initial configuration for simulations of both equilibrium configurations and dynamics.

### Equilibrium configurations and quasi-static changes

Equilibrium states were obtained by quenching the system from an initial state with tension fluctuations (Δ*T/T*_0_ = 0.5) to an equilibrium state (Δ*T/T*_0_ = 0) for each parameter set. Quasi-static changes of the relative adhesion at equilibrium where performed by first quenching the system to equilibrium at a given parameter value, and then performing a small change in the parameter (*W/T*_0_) and letting the system relax. Specifically, the system was initialized at a large value of the relative adhesion strength that ensured a confluent state (*W/T*_0_ = 1) and then quenched to a local equilibrium state. The relative adhesion strength was then progressively reduced to *W/T*_0_ = 0 by small changes (Δ*W/T*_0_ = −0.02). Similarly, the other branch was found by initializing the system at zero adhesion strength (*W/T*_0_ = 0) and progressively increasing it to *W/T*_0_ = 1 by small increments (Δ*W/T*_0_ = 0.02). After each adhesion adjustment, the system is relaxed to local equilibrium states (Supplementary Information).

### Introducing spaces between cells

Extracellular spaces are first introduced to initial configurations of confluent states as cells with different properties. Each vertex is replaced by a small triangular extracellular space centered around the original vertex position, and with the 3 new vertices located on the each of the original edges and of 1% their original length. These extracellular spaces then behave like ‘cells’ with different properties (see Methods above) and their size and geometrical features are determined by force balance at the vertices, as is the case for cells too. When two extracellular space ‘cells’ become neighbors, they are merged.

### Intermediate non-physical vertices

Intermediate vertices are introduced for both cell-cell contacts and free cell boundaries to allow for more realistic cell shapes. With intermediate vertices, individual edges consist of linear segments joined together to form a piecewise linear edge. The desired segment length is introduced as a parameter and the number of intermediate vertices for each edge is equal to the closest lower integer given by the ratio of instantaneous edge length to the segment length criterion. When the edge length increases (decreases), an intermediate vertex can be added (deleted) following the criterion just described. As the number of intermediate vertices changes, intermediate vertex positions are reassigned uniformly along the edge. If the longest segment is longer than twice the shortest segment in a given edge, intermediate vertex positions are also reassigned uniformly along the piecewise linear line.

### Treatment of topological transitions

T1 transitions occurs in our description when a given edge length is shorter than a critical length *ℓ*_*c*_ = 0.01*×*2*πL*_0_. When a T1 transition leads to the formation of a new vertex between 3 cells, we introduce a small triangular extracellular space ‘cell’ and let it evolve in time, as described above. Free cell boundaries (a boundary of a cell and the extracellular space) occasionally intersect each other due to the system dynamics. This event corresponds to the formation of a new contact between two cells. Therefore, when an intersection between any two free boundary edges is detected, a new cell-cell contact is introduced, splitting the extracellular space in two.

### Cell trajectories and mean squared displacement (MSD)

To obtain the Mean Squared Displacement (MSD) we first computed the cell centers (centroid of polygon), 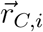, for each time point, *t*, based on vertex positions. Then, the cell trajectories were obtained by monitoring the changes of cell centers. Using the cell trajectories 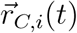, we obtained the MSD according to

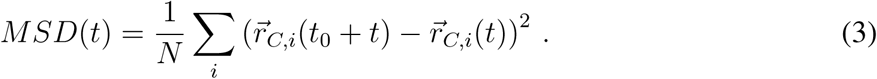

MSD values were averaged over all cells and also all initial time points *t*_0_ for any given set of model parameters.

### Application of a step strain

A large step strain was applied by deforming the simulation box from a square to a parallelogram of the target shear strain (150%). An affine deformation is applied to all vertex positions when imposing the step strain. Lees-Edwards periodic boundary conditions were imposed throughout the simulation to avoid mismatch of cell geometry across system boundaries.

### Stress calculation

The non-dimensional stress tensor can be computed from the transient tissue geometry with knowledge of junctional tensions and cell pressures ^39, 40^ (lowercase indicates normalized quantities; Supporting Information), namely

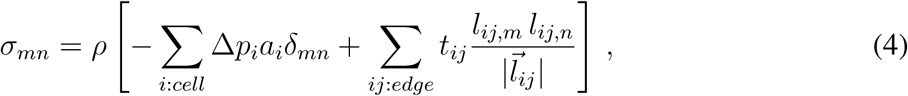

where Δ*p*_*i*_ ≡ *p*_*i*_ − *p*_0_, *m, n* are indices indicating the spatial direction (*m* = *x, y* and *n* = *x, y*), *a*_*i*_ is the dimensionless cell area and 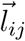 is the vector form of the edge length between cells *i* and *j*. The shear stress term can be written as

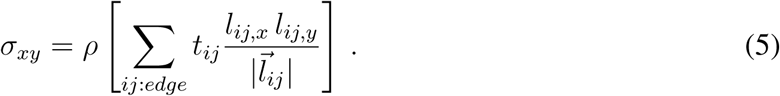

### Active shear stress

Due to tension fluctuations, the macroscopic shear stress shows fluctuations around zero. To quantify the magnitude of these active shear stress fluctuations, the shear stress values are monitored over a time interval of 10^3^ *τ*_*R*_ and their standard deviation is computed for each parameter set (no macroscopic imposed strain). The level of active shear stress corresponds to the computed standard deviation of shear stress, and increases approximately linearly with the magnitude of tension fluctuations for a given relative cell adhesion strength (Supplementary Information).

### Zebrafish husbandry, lines and experimental manipulations

Zebrafish (*Danio rerio*) were maintained under standard conditions^41^. The cdh2^tm101^ mutant line ^36^ was used to disrupt adhesions between cells, otherwise phenotypically wild-type lines were used. Animal husbandry and experiments were done according to protocols approved by the Institutional Animal Care and Use Committee (IACUC) at the University of California Santa Barbara. Transgenic lines Tg(hsp70:secP-mCherry)^p142^ and Tg(h2afva:eGFP)^kca6^ were used to visualize extracellular spaces and nuclei, respectively. Both Tg(bact2:mem-neonGreen-neonGreen)^hm40^ and Tg(actb2:memCherry2)^hm29^ transgenic lines were used to visualize cell membranes in different experiments. In some cases, membranes were labelled by injection with 80-100 pg membrane-GFP mRNA at 1-2 cell stage.

### Imaging

In all cases, 8-10 somite stage zebrafish embryos were mounted in 1% low-melting point agarose in a glass bottom petri dish (MatTek Corporation) for a dorsal view of the tailbud and imaged at 25°C using a 40x water immersion objective (LD C-Apochromat 1.1 W, Carl Zeiss) on an inverted Zeiss Laser Scanning Confocal (LSM 710, Carl Zeiss Inc.).

For kymograph of cell-cell contacts, confocal timelapse data of MPZ and PSM regions in Tg(h2afva:eGFP)^kca6^ x Tg(actb2:memCherry2)^hm29^ double transgenic embryos were acquired (2 second time intervals for 30 minutes). The intensity profile of a single junction was tracked over 300 seconds and measured over a 5-pixel width segment line to minimize the error induced by the estimation of the junction location.

Images of extracellular spaces were obtained using an outcross of Tg(hsp70:secP-mCherry)^p142^ and Tg(bact2:mem-neonGreen-neonGreen)^hm40^, to label the extracellular spaces and cell membranes, respectively. To trigger the expression of mCherry in the extracellular space, a 1 hour heat shock in a water bath at 39°C was performed when the embryos were at 75% epiboly stage. Confocal stacks through PSM and MPZ tissues were acquired with a z-step size of 0.55*µm*. Cell membranes were segmented using CellPose^43^. The resulting segmented image is overlaid on the confocal section showing both the secreted fluorescent reporter in the extracellular spaces and cell membranes (Fig. 5**b**).

### Tracking of cell movements in the PSM of N-cadherin mutants

Both mutant (*cdh2*^−/−^) and sibling (*cdh2*^+/−^) embryos were injected with 80-100 pg each of H2B-mRFP mRNA and membrane-GFP mRNA at 1-2 cell stage to label nuclei and membranes, respectively. Confocal stacks through the PSM were acquired with a z-step size of 1*µm* and time interval of 1 minute for 30 mins, and processed using Imaris (Bitplane). Data were smoothed using a 1-pixel gaussian filter, to correct for photobleaching over time the normalize timepoints function was used, then attenuation correction was applied to correct for z-attenuation. After processing, data were cropped to the tissue of interest, then nuclei were detected using the spots function, and tracked using the Brownian motion algorithm. Nuclei positions were output for further analyses. The Mean Squared Relative Displacement (MSRD) is calculated from the cell trajectories^13^. MSRD in 3D corresponds to 3 times the MSD in 2D.

### Measurement of cell shapes and shape factor

To measure cell shape and the shape factor we used Tg(h2afva:eGFP)^kca6^xTg(actb2:memCherry2)^hm29^ double transgenic embryos (wild type phenotype) and N-cadherin (*cdh2*^−/−^) mutants injected with 80-100pg H2B-mRFP mRNA and 80-100 pg membrane-GFP mRNA at 1-2 cell stage. In all cases, confocal stacks were acquired with a z-step size of 0.55 *µm*. Segmentation analysis of membrane labeled images were performed in FIJI ^44^. To measure cell shape, we chose the measurement plane of the confocal section when the nucleus area was maximized. The shape factor was calculated by fitting a polygon to the shape of the cell membrane signal. To visualize dynamics of cell shape contour, the template matching plugin^45^ was first applied to measured timelapses of membrane-labeled images to remove drift and an average intensity projection image was obtained from the drift corrected timelapse images.

## Data availability

All data is available from the authors upon reasonable request.

## Code availability

The code developed for this manuscript is available from the authors upon reasonable request.

